# A Hierarchical Anti-Hebbian Network Model for the Formation of Spatial Cells in Three-Dimensional Space

**DOI:** 10.1101/264366

**Authors:** Karthik Soman, Srinivasa Chakravarthy, Michael M. Yartsev

**Affiliations:** Bhupat and Jyoti Mehta School of Biosciences, Department of Biotechnology, Indian Institute of Technology Madras, Chennai, Tamilnadu, India - 600036; Department of Bioengineering and the Helen Wills Neuroscience Institute, University of California–Berkeley, Berkeley, CA 94708, USA

## Abstract

Three dimensional (3D) spatial cells in the mammalian hippocampalformationare believed to support the existence of 3D cognitive maps. Modeling studies are crucial to comprehend the neural principles governing the formation of these maps, yet to date very few have addressed this topic in 3D space. Here, we present a hierarchical network model for the formation of 3D spatial cells using anti-hebbian network. Built on empirical data, the model accounts for the natural emergence of 3D place, border and grid-cells as well as a new type of previously undescribed spatial cell type which we call plane cells. It further explains the plausible reason behind the place and grid-cell anisotropic coding that has been observed in rodents and the potential discrepancy with the predicted periodic coding during 3D volumetric navigation. Lastly, it provides evidence for the importance of unsupervised learning rules in guiding the formation of higher dimensional cognitive maps.

## INTRODUCTION

Empirical studies in rodents show that hippocampal and parahippocampal regions contain a multitude of ‘spatial cells’ that contribute to the creation of a cognitive map for navigation. Rodent hippocampus is reported to have ‘place cells’that fire at localized regions of the navigated space (Moser et al., 2014; O'Keefe and Dostrovsky, 1971). Medial entorhinal cortex (MEC) of rats is reported to contain ‘grid cells’ that activate when the animal passes through one of multiple locationsin space where the spatial alignment of these firing locations exhibits a strikingly hexagonal grid-like firing pattern (Hafting et al., 2005; Moser et al., 2014). Direction sensitive cells that encode the head direction of the animal in the yaw plane are reported from a wide range of regions including postsubiculum and MEC (Taube and Bassett, 2003; Taube et al., 1990a, b). Subiculum and MEC are reported to have ‘border cells’ that encode the borders of the environment (Bjerknes et al., 2014; Lever et al., 2009; Solstad et al., 2008).

Efforts to determine the precise coding for 3D space in rodents are ongoing, yet appeared to manifest in contradicting results under different behavioral conditions in which the behavior of the animal was constrained to different arrangements of two-dimensional (2D) planes (Bassett and Taube, 2001; Calton and Taube, 2005; Knierim and McNaughton, 2001; Stackman and Taube, 1998; Stackman et al., 2000). In parallel, results on 3D spatial maps have been obtained from bats, a mammal that naturally navigated in unconstrained fashion during flight while exhibiting spatial movement patterns that span the entire 3D volumetric space (Ulanovsky, 2011; Ulanovsky and Moss, 2007). Bat hippocampus is reported to contain place cells that are active in confined 3D volumes (Yartsev and Ulanovsky, 2013). 3D head direction (HD) cells, which form an internal compass for animal’s 3D navigation, have been reported in the dorsal presubiculum of the Egyptian fruit bats (Finkelstein et al., 2015). These HD cells code for the direction of motion in terms of the three Euler angles viz. azimuth, pitch and roll (Finkelstein et al., 2015). Grid cell activity has thus far only been reported from the MEC of bats during 2D navigation, yet has been shown to exhibit many of the classical grid cell features that have previously been reported in rodents, such as hexagonal firing fields and gradient in grid scale across the dorso-ventral MEC axis (Yartsev et al., 2011). Apart from pure grid cells, bat MEC is also reported to have other spatial cells viz. conjunctive grid cells, pure head direction cells and border cells (Yartsev et al., 2011) yet these have thus far also only been studied in 2D environments.

These rich empirical data raise difficult questions about spatial maps in higher dimensions: What is the learning rule for the formation of empirically reported 3D spatial cells? What form of symmetry does a grid cell take in higher dimensions? What contributes to the isotropic and anisotropic coding schemes of spatial cells and why different mammals differ from each other with respect to 3D spatial coding properties? Can there exist other kinds of spatial cells to represent the space in higher dimensions? A systematic comprehensive computational model is pertinent to decipher these queries. Although a corpus of computational models exists in the case of the 2D navigation problem (Burak and Fiete, 2009; Burgess et al., 2007; Burgess and O’Keefe, 2011; Bush and Burgess, 2014; Fuhs et al., 1998; Fuhs and Touretzky, 2006; Kropff and Treves, 2008; Soman et al., 2017; Widloski and Fiete, 2014; Zilli and Hasselmo, 2010), models of 3D navigation are conspicuous by their near total absence. Mathiset *al* treated the probable nature of grid like representations in higher dimension as a packing problem and concluded that the periodic grid like pattern in 3D navigation may take Face Centered Cubic(FCC) lattice structure since it has the highest packing ratio (Mathis et al., 2015). A rate adaptation network model, where the grid cell is assumed to receive place cell inputs - empirically validated in the case of 2D navigation in rodents (Bjerknes et al., 2014; Bonnevie et al., 2013), but not yet in bats nor in 3D navigation - suggests the possibility of an asymptotic state of FCC or Hexagonal Close Packing (HCP) lattice grid structure in 3D space (Stella and Treves, 2015). Apart from delineating the possible nature of grid fields in 3D space (which is still an open question), no other aforementioned experimental findings (such as 3D place and border cells) are addressed by any of these models. This sparsity of modeling efforts designed to address the encoding of 3D space in which most animals live in, represents a major challenge to the understanding of its underlying neural computations.

To bridge this major gap, we propose here a hierarchical network model which accounts for the formation of *all* spatial cells reported to-date in 3D space. The model shows how 3D spatial maps could be formed using unsupervised anti-hebbian neural network while the animal follows a naturalistic complex trajectory in 3D space. The presented generalized framework not only accounts for a gamut of empirical results in 3D navigation but also makes significant predictions on the learning rule, isotropy in spatial coding and possible nature of periodic grids fields in 3D. Lastly, it also makes direct predictions for the possible existence of other kinds of novel 3D spatial representations that have yet to be reported in animals navigating in 3D. To the best of our knowledge, this is the first hierarchical systems-level modeling approach taken to model 3D spatial cells.

## RESULTS

### Model architecture for 3D spatial cells

The proposed model is driven by the movement of the virtual animal during its active exploration in the 3D space. Simulated flight trajectory concurs with the empirical trajectory of bats in terms of its azimuth and pitch distribution statistics (Finkelstein et al., 2015; Yartsev and Ulanovsky, 2013). Azimuth angle is sampled from a uniform distribution which spans the entire 360° angular space (Figure 1B). However, since the animal’s flight is devoid of sharp dives and ascents, its pitch exhibits a narrower range than azimuth (Figure 1C).

**Figure 1:**
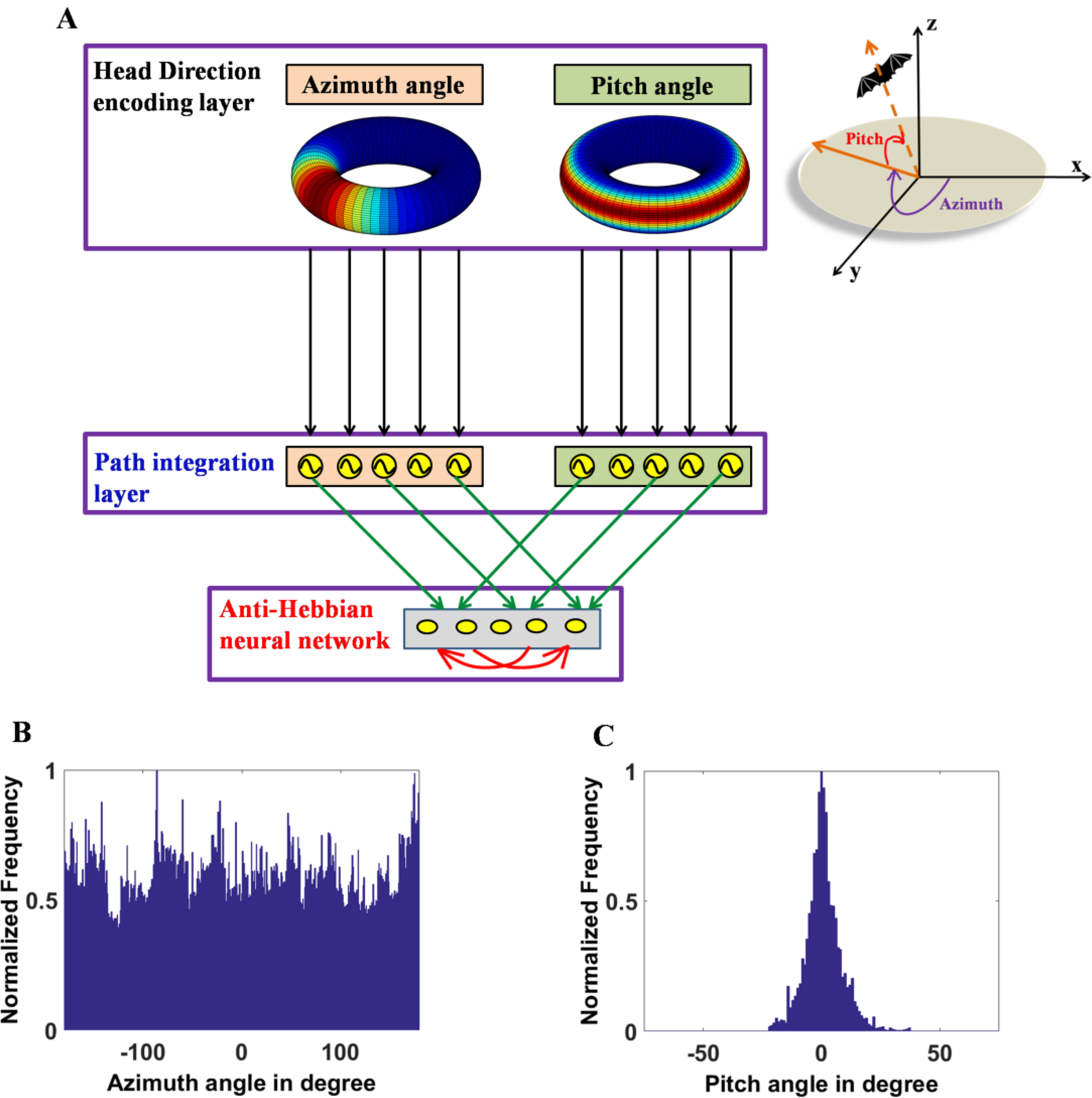
Model architecture and trajectory statistics. (A) The architecture of the proposed model. Illustration of the hierarchical nature of the model starting from the head direction encoding layer with parallel layers that code for the azimuth and pitch angle. Inside the head direction encoding layer, toroidal representations for pure azimuth (left) and pure pitch (right) are shown. Head direction layer ensures one-to-one connectivity with the corresponding oscillatory path integration layer which further converges to the anti-hebbian network. Green arrows indicate the afferent synaptic weight connections that are trainable by hebbian rule and the red arrows indicate the lateral inhibitory synaptic connections that are trainable by anti-hebbian rule. Top right side to the figure shows the depiction of the pitch and the azimuth angle in the 3D Cartesian coordinate system. (B) Angular distribution of the azimuth direction of the simulated trajectory. Azimuth angle of the simulated trajectory is uniformly distributed. (C) Angular distribution of the pitch direction of the simulated trajectory. Pitch angle of the simulated trajectory has a narrow range as shown in the figure with a Gaussian distribution (*μ* = 0, *σ^2^* = 58.25).

The model adopts a hierarchical architecture comprising of: Head Direction (HD) encoding layer, Path integration layer and finally Lateral Anti-Hebbian Neural (LAHN) layer (Figure 1A). Empirical studies on bats report the existence of separate neural ensembles to code for the three Eulerian angles-azimuth, pitch and roll, with the roll-coding population being the smallest (Finkelstein et al., 2015). Conforming to this empirical result, HD layer in the proposed model has two parallel layers viz. Azimuth and Pitch direction encoding layers. Roll encoding cells are neglected in the model, since Egyptian fruit bats (the only bat species to date from which hippocampal neural data has been recorded during flight conditions) seldom use roll direction for navigation (rather more so for landing) and also empirically roll coding population is insignificant (Finkelstein et al., 2015). Since azimuth cells are higher in distribution compared to pitch cells (Finkelstein et al., 2015), they are taken in a ratio of 7:3 in the model, as reported empirically in the bat (Finkelstein et al., 2015). Irrespective of this distribution, the preferred directions of both cell types span the complete 360° angular space. Hence, in the model, these cell types compute the projection of the current heading direction (azimuth or pitch depending on the cell type) on to its preferred direction and forward pass the information to the downstream path integration layer. Like HD layer, path integration layer also splits into two parallel layers of azimuth and pitch respectively and it ensures one-to-one connectivity with the upstream HD layer. It is an array of phase oscillators with low frequency compatible with reports in bats (0.5 Hz, see(Heys et al., 2013; Yartsev and Ulanovsky, 2013; Yartsev et al., 2011)). It achieves path integration by integrating the afferent encoded direction and speed information of the animal’s flight into the phase of the oscillator.

In the final hierarchy of the model, the information from the parallel pathways (azimuth and pitch pathways) converges together and form afferent input to the anti-hebbian network. Hence LAHN receives the integrated information of the 3D space in which the animal navigates. LAHN is a recurrent neural network whose afferent weight connections are updated using hebbian rule i.e. higher correlation of pre and post synaptic neural activity strengthens the synaptic weights between them (Barlow and Foldiak, 1989; Földiak, 1990). Lateral recurrent connections are updated using anti-hebbian rule i.e. higher correlation of pre and post synaptic neural activity suppresses the synaptic weights between them (Földiak, 1990). Anti-hebbian lateral connections induce competition among the neurons whereas the afferent hebbian connections enable extraction of principal components from the input data (Földiak, 1990; Oja, 1982; Sanger, 1989). An iterative learning network also has the advantage of analyzing the temporal evolution of the spatial cells. Owing to its generalized architecture and the local learning rules, LAHN qualifies as a biologically plausible neural network.

After training the model, the LAHN neural activity is tracked along realistic 3D flight trajectory of the animal. Whenever the activity of a neuron crosses a threshold value, we assign this threshold crossing moment an action potential, which allows us to compute the firing field of that particular neuron.

### Emergence of 3D Place fields

While training the LAHN, some neurons evolve firing fields that eventually occupy a confined volume in the 3D space (Figure 2A). Once the network converges, these simulated firing fields resemble the volumetric place fields reported from the flying Egyptian fruit bats (Figures 2B-2E). Place cells evolved from the model are further compared with their empirical counterparts with regard to their isotropic index. Place cells from the CA1 region of bat hippocampus are reported to have isotropic firing fields i.e. the firing fields show equal variance in all configurations of three orthogonal dimensions (Yartsev and Ulanovsky, 2013). This is quantified by fitting an ellipsoid to the 3D firing field and computing the ‘elongation index (*ξ*)’ as a measure of isotropy (Yartsev and Ulanovsky, 2013). *ξ* is the ratio of the largest to the smallest axis of the fitted ellipsoid. The Gaussian distribution fitted on the *ξ* (computed from 164 place cells obtained by retraining the LAHN tentimes) shows that a significant number of simulated neurons exhibit isotropic firing fields (mean = 1.3042, s.d = 0.2179) (Figure 2F).

**Figure 2:**
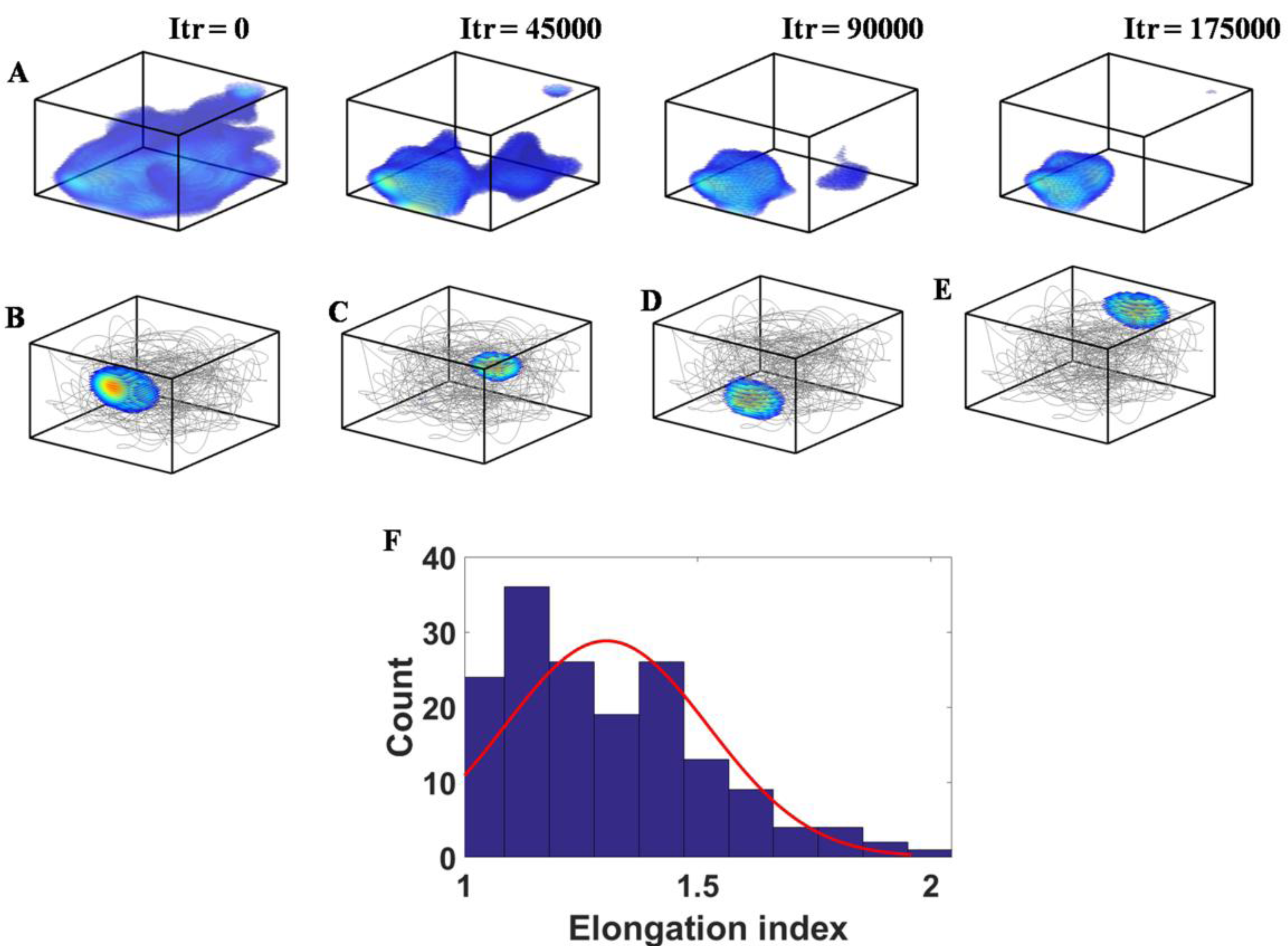
Emergence of 3D place cells and their isotropicnature. (A) Figure, from left to right, shows the snapshots of the temporal evolution of the localized volumetric 3D place cell firing field of a LAHN neuron that resembles the empirically reported 3D place cells. Iteration number is given above each box. (B)-(E) show the rate maps of four example 3D place cells from the trained model, overlaid on the flight trajectory traversed by the animal (gray lines). It is evident from the figure that the place cells code for different regions of the space. (F) The distribution of the elongation index of the fitted ellipsoids results in similar values to those observed empirically in freely flying bats (Yartsev and Ulanovsky, 2013). Elongation index is the descriptor of the isotropic nature of the firing map. The Gaussian curve fitted to the distribution (red curve) mainly shows the emergence of isotropic nature of place cells from the model (*μ*= 1.3042, s.d = 0.2179).

### Emergence of 3D spatial periodicity

In the bat experimental literature, head direction and place cells are the two spatial cells reported while the animal navigates in the 3D space (Finkelstein et al., 2015; Omer et al., 2018; Yartsev and Ulanovsky, 2013). Albeit grid cells have been reported in bats, it does not form a conclusive evidence for the existence of 3D grids because the neural activity was recorded while the animal was crawling on the ground (Yartsev et al., 2011). Although empirical studies to search for 3D grid cells have been performed in rodents (Hayman et al., 2011; Hayman et al., 2015), the existence of 3D periodic spatial representations is still unresolved yet predictions for its existence can be presented using the model, as we shall show next.

### Analysis of FCC symmetry in the simulated 3D grid fields

Apart from the spatially localized activity of the aforementioned place cells, LAHN neuronal ensemble shows spatially periodic activity too. Spatially periodic activity is vivid from the rate maps of the grid neurons (Figures 3A, 3C). However, autocorrelation maps are computed to analyze the symmetry of the grid periodicity in higher dimensions (Figures 3B, 3D). Previous modeling studies have predicted the possibility of Face Centered Cubic (FCC) lattice structure for the grids in the 3D space owing to its higher packing fraction and thereby efficient coding of position (Mathis et al., 2015; Stella and Treves, 2015).

**Figure 3:**
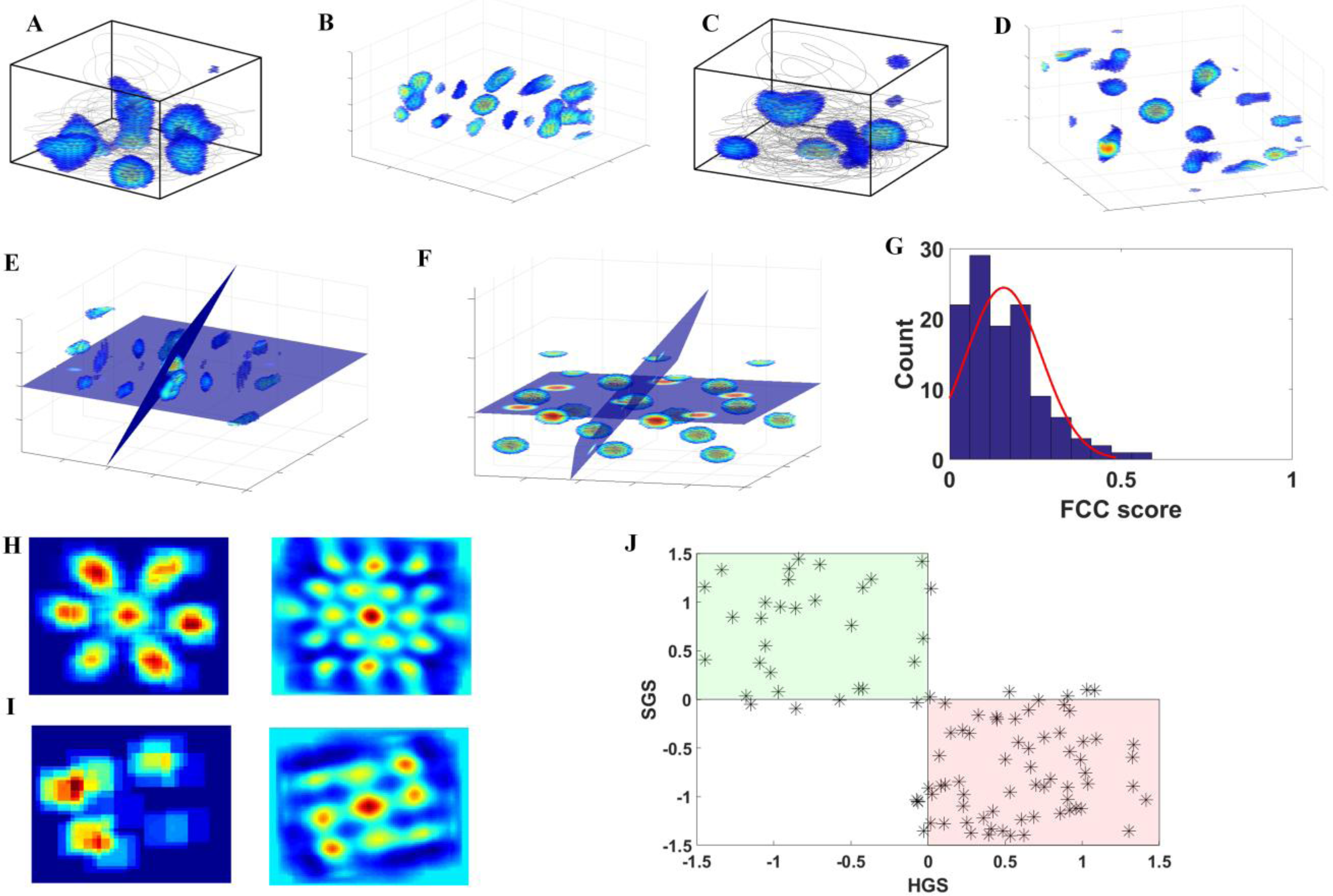
Emergence of 3D spatially periodic neurons and their symmetry. (A) 3D firing map of a LAHN neuron overlaid on the animal’s flight trajectory. (B) 3D autocorrelation map of the same neuron computed to observe the inherent periodicity of the spatial representation. (C) 3D firing map of another LAHN neuron overlaid on the animal’s flight trajectory. (D) 3D autocorrelation map of the same neuron computed to observe the inherent periodicity of the representation. Two neurons (A and C) are purposefully taken to show the difference in their periodic representations which will be made clearer through Figures H and I. (E) Two planes at 72° transecting through the central peak of the volumetric autocorrelation map of a spatially periodic LAHNneuron. This is done to check for any FCC symmetry in the 3D grid representation of that neuron. In the case of FCC symmetry, the transecting planesat 72° reveal hexagonal patterns on them. (F) Two planes at 72° transecting an analytically simulated FCC lattice structure through its central peak. This is done to compare with the previously mentioned transection and hence to compute the FCC score. (G) Distribution of the FCC scores of the spatially periodic neurons from the model. The Gaussian curve fitted to the distribution (red curve) shows that the neurons have less tendency to form the FCC symmetry *(μ* = 0.1575, s.d = 0.1098). (H) (Left) Projection of the 3D firing map shown in (A) onto the XY plane which shows hexagonal rate map (Right) Autocorrelation map of the 2D firing map to quantify the 60° periodicity (I) (Left) Projection of the 3D firing map shown in (C) onto the XY plane which shows square rate map (Right) Autocorrelation map of the 2D firing map to quantify the 90° periodicity (J) Scatter plot of the gridness score on 2D plane of HGS and SGS. Green shaded region shows the square grid area and the red shaded region shows the hexagonal grid area. The plot conveys that 60% of grids exhibit hexagonal symmetry and 26% exhibit square symmetry and remaining 14% show gridness scores at the border line whichcannot be used to discern between hexagonal and square symmetry.

FCC is a cubic lattice structure that results from stacking hexagonally arranged layers of spheres one above the other (Conway and Sloane, 2013). Considering three layers A, B, C where B and C are the translations of A, the sequence ABCABC results in an FCC structure (Conway and Sloane, 2013). In FCC symmetry, one can find four planes, at an angular difference of 72°, transecting through the center of FCC and each plane has peaks arranged in a hexagonal fashion (Stella and Treves, 2015). An ideal FCC lattice structure can be analytically simulated by a linear combination of four 3D waves whose wave vectors are at angles of 109.5° (Stella and Treves, 2015).

In FCC symmetry analysis of 3D grid fields, initially the autocorrelation map is transected into many slices that pass through the origin. Hexagonal gridness score (HGS) of each slice is computed. The slice with the highest gridness score is taken as the ‘reference plane’. Those planes at 72° from the reference plane are considered (Figure 3E shows two such planes) and an average of top three gridness scores of those planes is computed. A similar procedure is performed on the analytical FCC too (Figure 3F). An FCC score is computed by taking the ratio of these two average gridness scores (analytical FCC in the denominator). Negative average gridness score (if any) is set to zero so that FCC score will be between 0 and 1. Hence, if the spatial periodicity is close to FCC symmetry, the FCC score will be ~1. The aforementioned analysis on the LAHN spatially periodic neurons (114 spatially periodic neurons obtained by retraining the LAHN ten times) shows that the grid neurons from the model apparently do not show FCC symmetry (Gaussian distribution fitted on the FCC score has mean = 0.1575 and s.d = 0.1098) (Figure 3G).

Since the spatially periodic neurons from the model show less tendency towards the FCC structure, to check the symmetry adopted by the grids in the model, rate maps are projected on to the three orthogonal planes such as XY, YZ and XZ respectively (Figures 3H, 3I). 60° rotational symmetry is checked on each plane by computing their Hexagonal Gridness Score (HGS) (Hafting et al., 2005; Soman et al., 2017). Along with the HGS, Square Gridness Score (SGS) is also computed to consider the possibility of 90° planar symmetry (Soman et al., 2017). A positive gridness score on any one of the planes means that the grid cell shows a planar symmetry in the 3D space. Scattering the computed gridness scores on a 2D space of HGS and SGS (Figure 3J) confirmed the planar symmetry of the spatially periodic neurons from the model.

### Anisotropy in grid and place-cells coding on vertical plane

In the experimental study, when the rat forages on a 2D plane (XY), place-cells exhibit isotropic localized firing fields and grid-cells exhibit hexagonal representations. Yet, when the rat is made to climb over a pegboard (vertical YZ plane), the spatial representations of the same cells are reported to transform into an anisotropic representation, manifested as a single stripe for most place cells and multiple stripesfor the grid-cells (Hayman et al., 2011).This intriguing transformation exhibited by spatial neurons is still an enigma. This enigma is further enhanced by the empirical data obtained in bats. While bats place and grid-cells have exhibited similar firing patterns to those of rodents when the bats were navigating in 2D environments, it was found that the 3D firing patterns of place-cells in flying bats were dramatically different and in fact the complete opposite of the observation in rodents, i.e., they were isotropic. The proposed model is used to delve into this problem of anisotropy with the impetus of deciphering the reason behind this transformation. Specifically, we postulate that because the movement patterns of rodents and bats on 2D planes are far more similar to each other than in 3D space, that these differences in spatial movement patterns might lie at the core of the differences in neural coding. Such notions have been raised before, yet never actually tested.

To achieve this, two different trajectories are simulated on two different planes, one on horizontal and the other one on vertical plane (Figures 4 A, F). Since thus far only place-cells have been recorded in both species we begin our investigation there. When the virtual animal navigates on the horizontal XY plane (Figure 4A), the model gives rise to localized isotropic place firing field (Figures 4B, 4C) akin to the classical place cells. When the animal is made to navigate on the vertical YZ plane (Figure 4F), the previously formed isotropic firing field stretches out and forms stripe like firing field (Figures 4G, 4H). Grid neurons in the network exhibit hexagonal firing fields when the virtual animal navigates on the horizontal XY plane (Figure 4D). When the plane of navigation is switched from horizontal to vertical, it results in a transformation of the firing fields from hexagons (Figure 4D) to multiple stripes (Figure 4I). These symmetries are also reflected in the autocorrelation maps of the respective firing fields (Figures 4E,4J).

**Figure 4:**
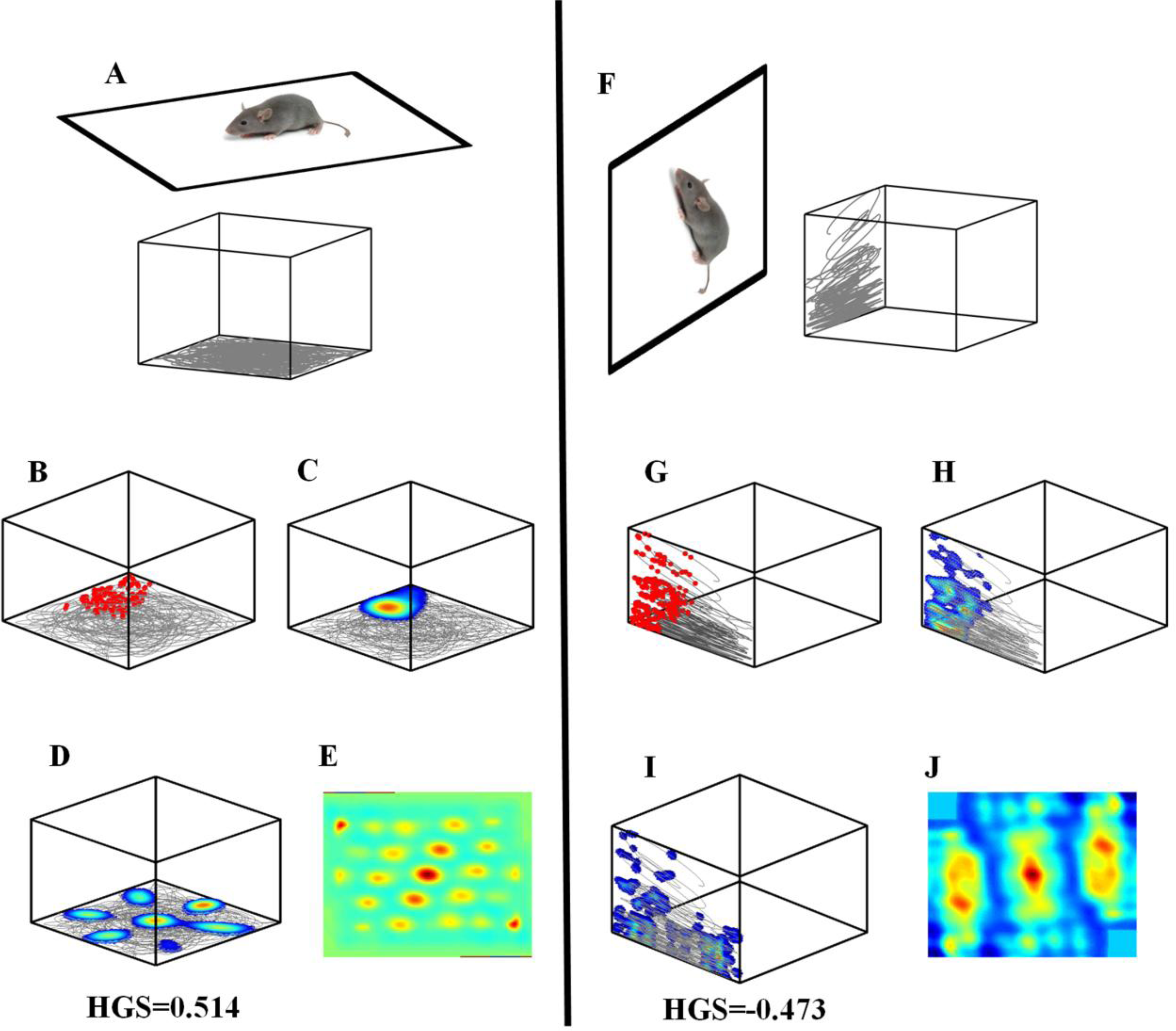
Horizontal and vertical anisotropy in grid cell coding. (A) A cartoon of a virtual animal navigating on horizontal XY plane (top) and the trajectory traversed by the animal (bottom) (B-C) Raw data (movement trajectories in gray and action potentials in red) and firing rate map of a LAHN place neuron in the model showing localized isotropic representation on the horizontal XY plane (D-E) Firing rate and autocorrelation map of a LAHN grid neuron in the model showing hexagonal representations on the horizontal XY plane (HGS>0). Hexagonal periodicity is vivid from the autocorrelation map and the gridness score (HGS). (F) A cartoon of a virtual animal navigating on vertical YZ plane (left) and the trajectory traversed by the animal (right) (G-H) Raw data (movement trajectories in gray and action potentials in red) and firing rate map of the same LAHN place neuron in the model showing stripe like representations on the vertical YZ plane. (I-J) Rate map data (movement trajectories in gray) and autocorrelation map of the same LAHN grid neuron in the model showing stripe like representations on the vertical YZ plane (HGS<0). Stripe periodicity is evident from the autocorrelation map.

Emergence of spatial representations in the proposed model isa result of the projection of path integration values on the weight vectors that maximize the variance of the output. It is important to emphasize here that the animal is not retrained on the vertical plane i.e. it is the same LAHN weight vectors (which is trained on the horizontal navigation) that are used for the navigation on both planes and exhibit different coding schemes. Hence, we hypothesize that the absence of hexagonal representations on the vertical dimension may not be a network property and could be attributed to the lesser variancein the vertical pitch distribution (Figure 1C). If this is the case, change in the variance of pitch distribution should reflect a corresponding change in the grid cell anisotropy too.

To test this hypothesis, three different trajectories with different pitch variancesare considered such as: trajectory with restricted motion on the vertical plane (Figure 5B) due to highly skewed pitch distribution (Figure 5A), trajectory with less restricted motion on the vertical plane (Figure 5F) due to an increase in the pitch variance compared to the first one (Figure 5E) and trajectory with a complete leverage to take any direction on the vertical plane (Figure 5J) due to uniform pitch distribution (Figure 5I). Since both grid and place neurons in the LAHN undergo similar stripe like transformation, the proposed hypothesis is analyzed only on one type of spatial cell i.e. grid cell (grid representation can also be quantified using the gridness score). LAHN grid neurons exhibit spatial periodicity on the vertical plane in all the three cases (Figures 5C, 5G, 5K), but the neural representations are different for each case. As the pitch variance increases, the spatial representations on the vertical plane gradually transform from stripes to hexagons which is evident from the respective neural firing fields (Figures 5C, 5G, 5K) and the autocorrelation maps (Figures 5D, 5H, 5L).

**Figure 5:**
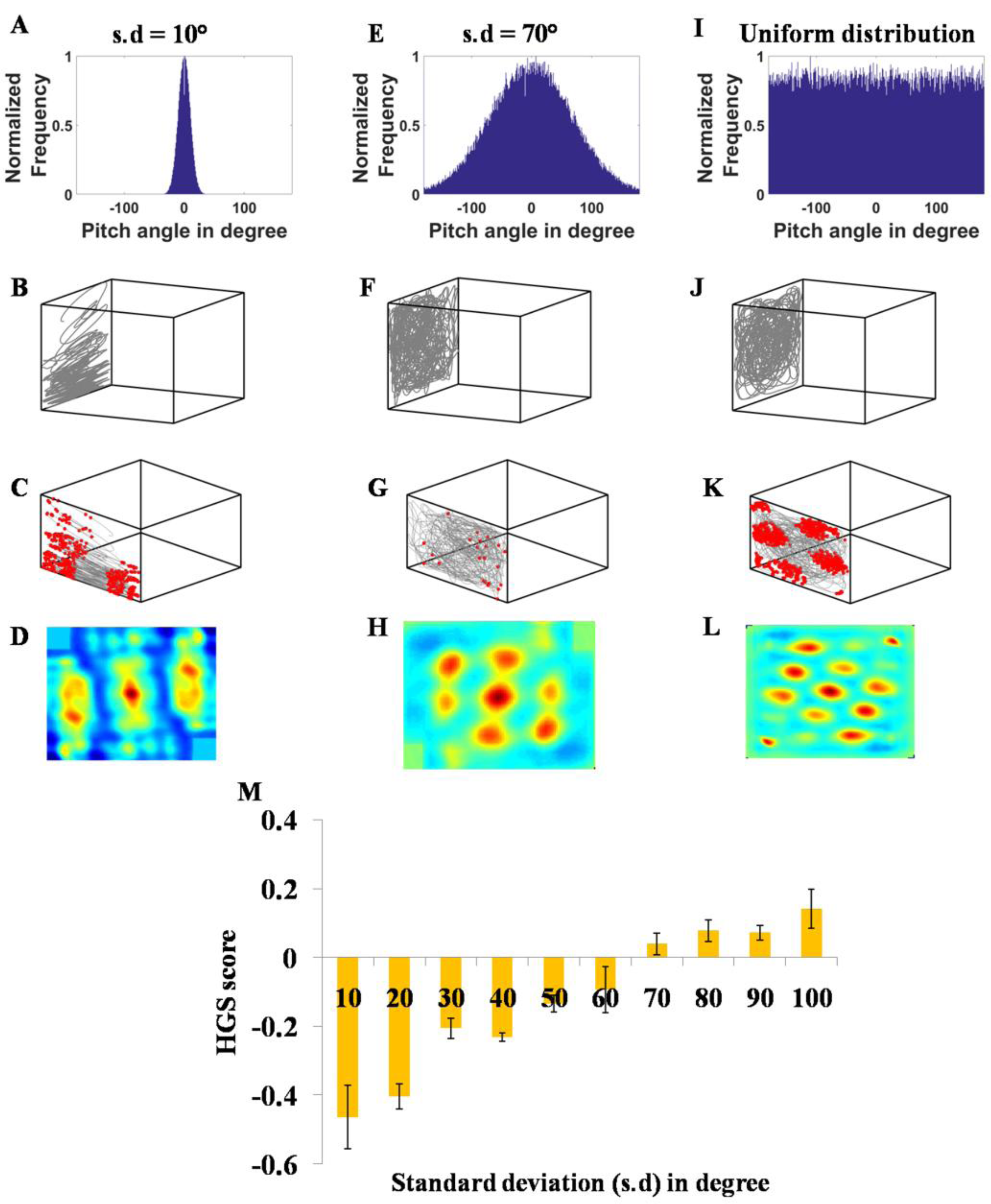
Influence of pitch variance on grid cell anisotropy. (A) Skewed pitch distribution with *μ =* 0; s.d = 10° (B) Trajectory simulated on the vertical wall using the skewed pitch distribution as shown in (A). (C) Stripe like firing fields of the LAHN neuron formed on the simulated vertical plane trajectory (D) Autocorrelation map of the representation that clearly shows the stripe periodicity (E) Less skewed pitch distribution with *μ* = 0; s.d = 70° (F) Trajectory simulated using less skewed pitch distribution. (G) Firing fields of the same neuron formed on the vertical plane trajectory. Loss of stripe like representation is vivid from the firing field. (H) Autocorrelation map of the representation that shows 60° grid symmetry. (I) Uniformly distributed pitch distribution. (J) Trajectory simulated using the uniform pitch distribution that gives the animal the leverage to choose any direction on the vertical plane. (K) Hexagonal firing fields of the neuron on the vertical plane trajectory formed from uniform pitch distribution. (L) Autocorrelation map of the same firing fields that shows cogent hexagonal grids. (M) Dependence of the 60° rotational symmetry of the neural representation formed on the vertical plane (quantified as HGS on the y axis) to the s.d of the pitch distribution of the animal’s trajectory on the same vertical plane. A flip of HGS from negative to positive is evident at 70° s.d (critical angle).

To obtain a conclusive resulton the relation between the pitch variance and the grid formation on the vertical plane, pitch distributions with standard deviation (s.d) ranging from 10° to 100° (with a step size of 10°) are considered. Average HGS of spatial representations formed on trajectories from each pitch distribution is plotted against the respective s.d (Figure 5M). A flip in the HGS score takes place at 70° s.d (Figure 5M). We call this as the *critical angle* at which the stripe representations get transform to the hexagonal representations. This aspect is addressed again in the discussion section.

### Emergence of 3D Border cells and plane cells

Border neurons convey information about the borders of the environment in which the animal navigates. These types of cells are reported from the hippocampal-parahippocampal areas of rodents like MEC and subiculum (Bjerknes et al., 2014; Lever et al., 2009; Solstad et al., 2008). This is also reported from 3D space exploring animals like bats (Yartsev et al., 2011). However, like 3D grid cells, the description of 3D border cells has yet to be provided empirically as these are thus far only reported in animals moving on 2D planes (Bjerknes et al., 2014; Lever et al., 2009; Solstad et al., 2008; Yartsev et al., 2011). Here, we report the possibility of the existence of border like activity even in higher dimensions. After training the model, some LAHN neurons start to exhibit higher activity near the borders of the environment (Figures 6A-6D). This border related activity is quantified by calculating the border score (BS) of the respective neuron on each orthogonal plane. If the BS on any two orthogonal planes exceeds a threshold value (the threshold value is chosen based on the previous empirical literature (Solstad et al., 2008)), then the neuron is classified as a 3D border cell. To compute the BS, we adopted the same approach as in the case of 2D navigation i.e. the ratio of the difference between the extent of a single field on any wall and the average distance to the nearest wall of each bin in the rate map (weighted by its activity) to the sum of these quantities (Bjerknes et al., 2014; Diehl et al., 2017; Solstad et al., 2008). Hence the model predicts the possibility of the existence of border cells during volumetric 3D navigation.

**Figure 6:**
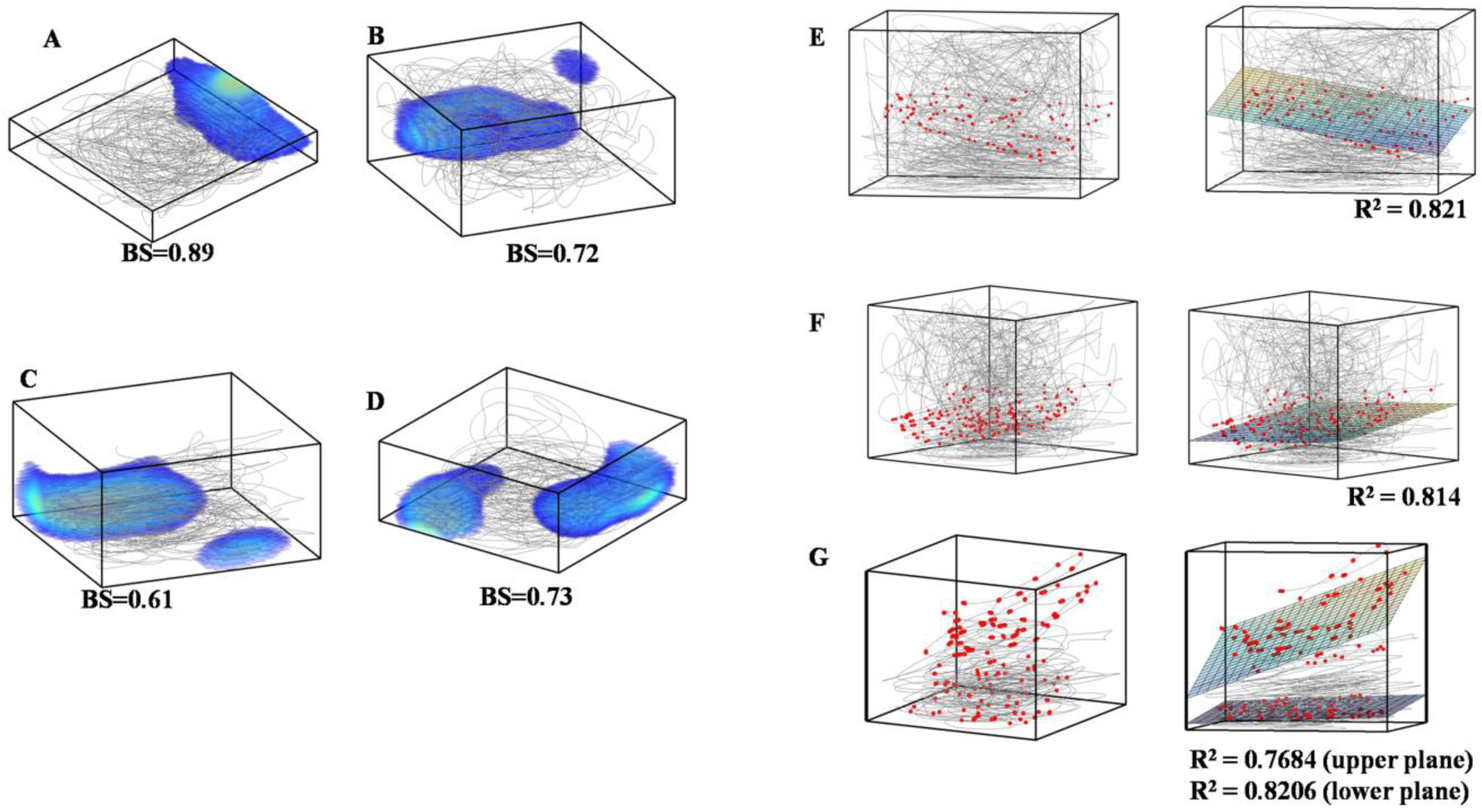
Emergence of 3D border and 2D plane cells. (A)-(D) 3D rate maps of four LAHN neurons that show affinity towards the 3D borders or walls of the box. Each 3D border cell has its own preferred border (either single or double preferred borders) to fire akin to the 2D border cells in the rodents. Border Score (BS) (average of top two BS) of each neuron is shown near to each box. (E)-(F) (Left) Raw data (movement trajectories in gray and action potentials in red) of three neurons that activate predominantly on a 2D plane in 3D space i.e. they are coding for a lower dimensional subspace and hence termed as plane cells (Right) 3D plane fitted to each plane cell firing field to quantify their ‘planeness’ in terms of the R^2^ value of the fitted plane. (G) (Left) Raw data (movement trajectories in gray and action potentials in red) of aneuron showing activity on multiple 2D planes and termed as stack cells. (Right) Two 3D planes fitted to the upper and lower firing fields. R^2^ value of each plane is shown near to the box.

Our model also revealed the possibility of a previously undescribed spatial cell type in 3D space which codes for 2D surface while the animal performs volumetric3D navigation. We call these neurons ‘plane cells’. Some LAHN neurons show this type of coding where neuronal firing fields are arranged on a 2D plane in a 3D space (Figures 6E-6G (left)). The amount of ‘planeness’ of a cell is quantified by the R^2^value of the plane fitted to the 3D firing field (Figures 6E-6G (right)). R^2^ value, which is the proportion of the variance in the dependent variable predictable from the independent variable, ranges from 0 to 1. The border cells also can have high plane index since it is also coding for a plane (border), or an entire face of the navigation volume. Hence for a cell to qualify as a pure *plane cell* it should have high R^2^ value (> 0.75) and less BS value (< 0.5). Apart from the plane cells which have a single plane of firing field, we also observed LAHN neurons with multiple planes of firing fields (Figure 6G) and termed them as *stack cells.* We believe that this model prediction on the possible existence of plane cells or stack cells is critical because coding of lower dimensional subspace (a plane) while performing navigation in higher dimensions (3D space) may be functionally relevant for animals navigating in 3D volumetric environments. For instance, a horizontal plane cell can convey the information about the altitude of the animal’s flight from the ground which is critical for the efficacy of its navigation.

### Distribution of cell types from the model and the influence of network size on spatial cells

Based on indices such as spatial information (SI), grid score (GS), border score (BS), plane index (PI), a quantitative analysis is done to account for the distribution of cell types that emerge from the model. Initially the distribution of spatial and non-spatial cells in the model is computed based on the SI index of each LAHN neuron. Based on the SI index, 95% of LAHN neurons evolve as spatial cells (based on average SI obtained after retraining the LAHN for twenty times) and remaining 5% as non-spatial cells (Figure 7A). This means that the network is able to code for one or the other spatial variables like place, grid, border, plane etc. Further analysis is done to check the distribution of the spatial cells formed from the model. The spatial cells are clustered in the spatial descriptor space based on their threshold values (Figure 7B). The distribution values convey that, out of the 95% spatial cells formed from the model 32.43% are place cells, 23.97% are grid cells, 28.1% are border cells and 15.5% are plane cells (Figure 7C).

**Figure 7:**
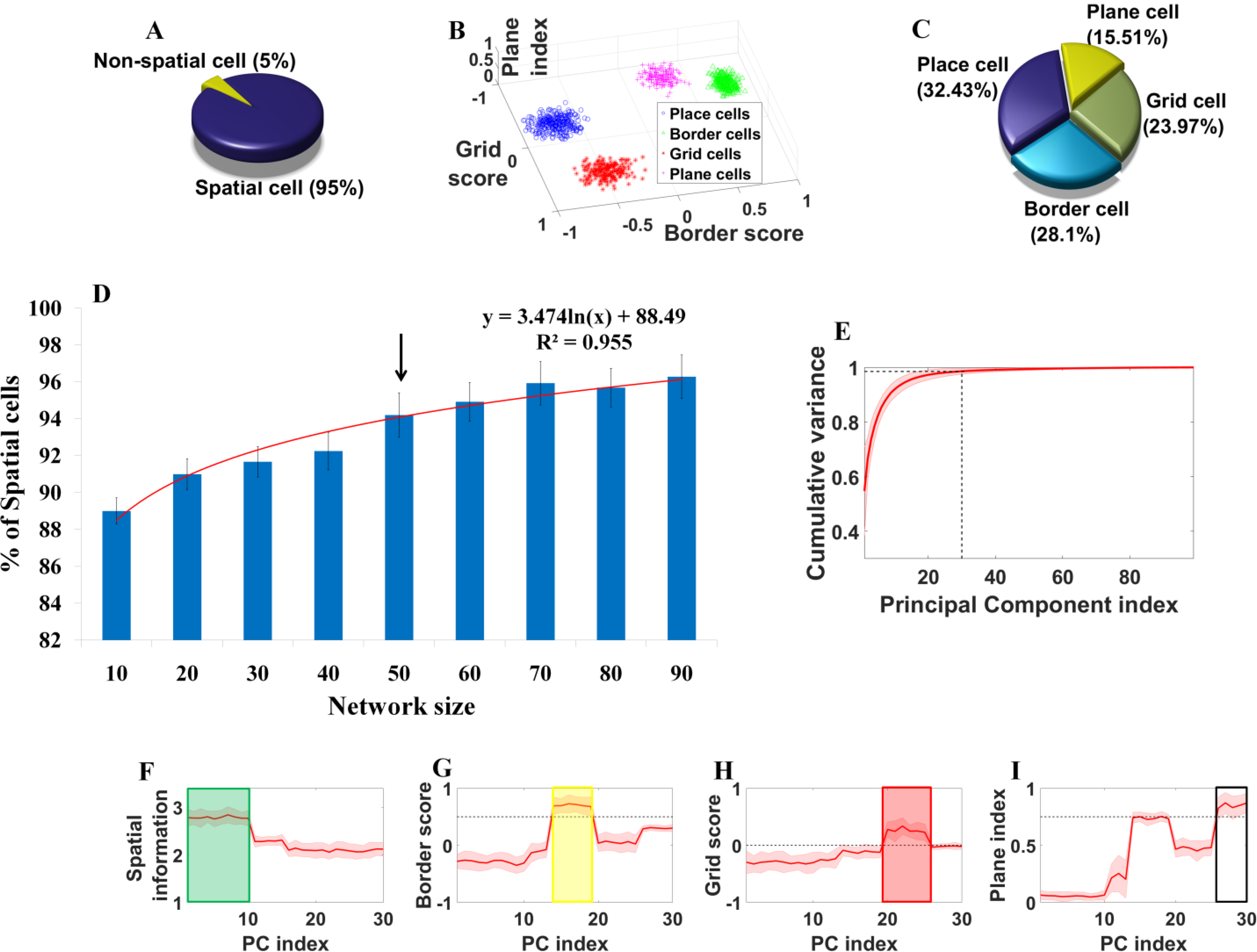
Distribution of cell types and the influence of network size on the model. (A) The Pie chart shows the distribution of spatial cells and non-spatial cells from LAHN network with 50 neurons. Cells are classified as spatial cells based on their spatial information (SI) index. Values are averaged over 20 trials (i.e. training the LAHN 20 times). Distribution shows that most of the neurons in the network after training code for one or the other spatial variable. (B) Clustering of different spatial cell types into place cells, border cells, grid cells and plane cells in the descriptor space (grid score, border score and plane index) (C) Pie chart shows the distribution of different spatial cell types from LAHN network with 50 neurons. Distribution shows that out of the 95% of spatial cells formed, 32.43% emerge as place cells, 23.97% emerge as grid cells, 28.1% form border cells and 15.5% form plain cells. (D) Graph (mean±s.d) shows the influence of LAHN size on the distribution of spatial cells. The logarithmic trend line (red curve) shows that the percentageof average spatial cell distribution increases as the size of the network increases. The black arrow indicates the network size from which the calculations in panels A-C are drawn. (E) Shaded plot of the cumulative variance graph of the principal components. Intersection of the black dotted lines shows the coordinate points at which the cumulative variance reaches 99% at 30^th^ principal component. (F) (I) Variation of spatial cell descriptors such as spatial information index, border score, grid score and plane index respectively with respect to the principal component (PC) index (For e.g. PC index 1 represents the first principal component and so on and so forth).

Network size is a critical factor in the present modeling study. Previous modeling works on grid cells in 2D environments have also shown the criticality of this factor (Burak and Fiete, 2009). Hence, a range of network sizes (10 to 90 neurons with a step size of 10) are considered and each network is trained twenty times to check the percentage of spatial cells formed for each time. As in the previous case, spatial cells are defined based on their SI index. Here, we do not account for the explicit spatial cell types instead we are interested in the influence of network size on capturing any spatial variable. The trend line shows that as the size of the network increases it accounts for more spatial features and form more spatial cells (Figure7D). However, there is an upper bound on the size of the network that can be used in the current model. This is because of the learning rule of LAHN. It has been previously proven that if the afferent and the lateral connections of a network are trained using hebbian and anti-hebbian learning rules respectively, weights in such a network convergeto the subspace spanned by the principal components of the input data (Barlow and Foldiak, 1989; Földiak, 1990). Hence LAHN can be considered as a neural network implementation of Principal Component Analysis (PCA). PCA is an orthogonal linear transformation that rotates the data to the maximal variance direction and hence performs dimensionality reduction of the input with minimal loss of information (Wold et al., 1987). Taking this into consideration, LAHN represents the path integration inputs with minimal number of neurons where each neuron specializes to code for some unique feature of the space and eventually turns out into one of the spatial cell types as shown above. Hence, the maximum network size that can be implemented using LAHN is *N-1* where *N* is the total number of path integration neurons. If the network size is more than N 1, the network carries redundant information which is not an optimal way of coding any stimulus.

### Formation of spatial cell bands on the eigen spectrum

The LAHN network has been previously shown as the neurally plausible implementation of Principal Component Analysis (PCA) (Barlow and Foldiak, 1989; Földiak, 1990). PCA is an orthogonal linear transformation that rotates the input to that direction which maximizes its variance and has been widely used for reducing the dimensionality of the input data with minimal loss of information (Wold et al., 1987). Here, the analysis of LAHN network shows that for a network size of 50 neurons, around 95% cells extract some spatial features and qualify themselves as spatial cells by increasing their SI index (Figure 7A). The spatial cell types which we consider here such as place cells, border cells, grid cells and plane cells show a defined way of distribution among themselves (Figure 7C). To analyze the reason behind this, a direct PCA is done on the input path integration matrix. The cumulative variance graph shows that the first 30 principal components (PC) (out of 100) capture ~99% of the input variance (Figure 7E). Hence further analysis is done only using the first 30 PCs. Spatial descriptors such as spatial information, border score, grid score and plane index are computed for the rate maps obtained from the projections of the path integration values on to each principal component. Graphical plot of each descriptor (averaged over 35 trajectories) with respect to the principal component index (upto first 30 PCs) shows a clear band of spatial cell distribution in the eigen-spectrum (Figures 7F-7I). Place cells that show high spatial information index (because of their localized activity) show a band in the high variance side of the eigen-spectrum (green box in Figure 7F). Border cells whose border scores are greater than the border threshold occupy the middle band of the eigen-spectrum (yellow box in Figure 7G). Grid cells that show positive grid scores occupy their position next to the border cell band in the eigen-spectrum (red box in Figure 7H). Plane cells whose plane index cross the plain threshold come next in the eigen spectrum (white box in Figure 7I). Hence arranging the spatial cell bands in the descending order of their principal component variance goes as: place cells, border cells, grid cells and plain cells. This ordering in the eigen-spectrum shows correlation with the spatial cell distribution from the LAHN (Figure 7C). This shows a close relation with the spatial map formation and PCA-like learning rule.

## DISCUSSION

### Summary

We have presented here a hierarchical network model for the formation of a spectrum of spatial cells in the 3D space. The model captures empirically reported results such as the emergence of 3D place cells (reported from Egyptian fruit bats (Yartsev and Ulanovsky, 2013; Yartsev et al., 2011)), isotropic nature of 3D place cells during volumetric navigation(Ulanovsky, 2011; Yartsev and Ulanovsky, 2013) and the anisotropy in grid cell coding on vertical plane as reported in rodents (Hayman et al., 2011). The model also accounts for the puzzling anisotropy of grid-cell coding as reported in rodents by revealing the relationship between the anisotropy and the variance in the pitch distribution of the animal’s trajectory. While 3D spatial cells types such as grid-cells and border-cells have yet to be reported in an animal navigating in the complete volumetric 3D space, the model makes explicit predictions about their expected properties: firstly on the spatially periodic representations in 3D space the model contests the emergent periodic structure with the FCC symmetry and shows the possibility of a planar symmetry in 3D space rather than FCC like lattice structure. This is in partial agreement with the empirical reports that support planar symmetry of the spatial representations in 3D space (Hayman et al., 2015), albeit these report emerges from rodents that do not span the complete volumetric space during navigation. The model also predicts the existence of 3D border cells by showing the emergence of such cells whose 3D rate maps show bias towards any one or two borders (walls) of the box. Furthermore, the model gives rise to predictions for the existence of novel spatial cell types that have yet to be reported yet naturally emerge from this biologically plausible model. In detail, apart from the border cells, the model also shows the emergence of neurons that code for a lower dimensional subspace like a plane (plane cells) or more than one plane (stack cells). The aforementioned results are obtained by using anti-hebbian network which has an iterative way of learning its afferent synaptic weights using hebbian rule and the neurons interact among themselves through lateral inhibitory connections which are updated using anti-hebbian rule. These learning rules qualify the biological plausibility of the network and also strengthen the model predictions. To our knowledge, this is the first modeling attempt to explain the formation of all 3D spatial cells at a systems level by considering the current empirical results in 3D spatial navigation.

### Empirical results captured by the model

An extensive empirical study on the navigational behavior of the Egyptian fruit bats resulted in the discovery of 3D place cells in their dorsal hippocampus (Geva-Sagiv et al., 2016; Yartsev and Ulanovsky, 2013). This extends the implications of cognitive spatial maps onto the navigation in higher dimensional space. 3D place cells exhibit volumetric firing fields on their 3D flight trajectory and cover the entire space uniformly. Around 33% of Lateral anti-hebbian network (LAHN) neurons in the model show the emergence of localized firing fields akin to the aforementioned empirically reported 3D place cells (Figures 2B-2E, Figure 7C). Isotropic coding is an important feature that singles out 3D place cells of bats from their rodent counterpart (Yartsev and Ulanovsky, 2013). Place neurons from the model also exhibit isotropic 3D spatial encoding which is evident from the distribution of the elongation index obtained from a fitted ellipsoid (Figure 2F). The reason for isotropy has been attributed to the evolutionary pressure to encode and decode the 3D spatial information in an efficient manner (Yartsev and Ulanovsky, 2013). This seems to be a valid reasoning for the observed phenomena. However, we have proposed an alternative solution for the isotropic coding scheme manifested by the spatial cells as explained below.

As mentioned in the results section, place and grid cells are empirically reported to exhibit an intriguing transformation of their representations from classical localized isotropic place field and hexagonal grid firing fields to stripe like firing fields as navigation changes from horizontal XY to vertical YZ plane respectively (Hayman et al., 2011). In the model this happens without retraining the network i.e. the same trained network switches its coding schema once the navigation plane changes. If the same network connections show two different spatial coding schemes on different planes (Figures 4C, 4H and 4D, 4I), it means that the reason for this transformation is not a network property; rather it is a behavioral property or a contextual property. This is quite a significant result since it is generally believed that the type of a spatial cell depends on the model parameters alone. However, our modeling approach suggests, it is a joint effect of both the model parameters (weights) and the nature of the trajectory.Hence, we attribute this anisotropic coding schema to the variance in the animal’s direction distribution. Learning rules of the anti-hebbian network performs PCA like transformation i.e. projecting the high dimensional spatial inputs provided by the path integration oscillators to those weight vectors in the direction of maximal variance. Place and grid representationsare high level spatial encoding produced by combing the high variance-high dimensional sensory information. The less variance introduced into the vertical direction component (pitch) induces an anisotropic transformation to both place and grid cells in the model which is actually reflecting a variance change in the afferent inputs to these cells. Hence this means that the representations formed by the brain to encode any sensory stimulus may not be a static one; rather it could be a dynamic one.

### Predictions from the model

Empirical data on a strict 3D spatial navigation (i.e. an animal like bat making a flight in complete 3D space) is reduced compared to the enormous 2D navigation empirical data from the rodents (Barry et al., 2007; Bjerknes et al., 2014; Bonnevie et al., 2013; Diehl et al., 2017; Hafting et al., 2005; Lever et al., 2009; Moser et al., 2014; O'Keefe and Dostrovsky, 1971; Solstad et al., 2008; Taube and Bassett, 2003). Hence predictions on 3D spatial representations are very important for biologists to design their experimental protocol and to have an intuitive idea on what to look for. The proposed model makes many predictions on the 3D spatial representations.

Spatially periodic neurons like grid cells have been observed in crawling bats (Yartsev et al., 2011). There are also experimental efforts using rodents to study the 3D structure of grid cells (Hayman et al., 2011; Hayman et al., 2015). Hitherto, there are no conclusive results regarding the natures of 3D grid cells. In the proposed model, ~ 24% of LAHN neurons exhibit spatial periodicity (Figure 7C). The spatially periodic neurons are analyzed to check for any FCC symmetry which has been previously predicted as the possible form for the 3D grid structure on the basis of their optimal packing efficiency (Mathis et al., 2015; Stella and Treves, 2015). However, grid neurons in the model apparently do not exhibit FCC symmetry in its periodicity (Figure 3G). Further analysis of rotational symmetry on 2D orthogonal planes (Figure 3H, 3I) point to both hexagonal and square planar symmetrical nature of the grid neurons in 3D space. There are also empirical reports from rodent studies that favor the planar structure of grid cells in 3D space. However, it is very unlikely for the brain to choose a non-optimal structure (2D hexagons in 3D space) when there is an option to choose for an optimal structure (FCC lattice) in terms of the packing fraction. The question is what does a network like LAHN do when there is no enough variance/information regarding the 3D component because of highly skewed pitch distribution? In such a case, it may rely upon the very next option, a planar hexagon, which is an optimal structure in the subspace (2D space) which has high variance in the azimuth distribution. An increase in the pitch variance could possibly bring a transformation from planar to lattice symmetry. This need not be a simple transformation from planar to lattice; rather the grid cell could pass through other intermediate symmetries before reaching the optimal one. The possibility of this transition has been shown in a previous modeling study by Stella and Treves (2015) (Stella and Treves, 2015) and future studies should conduct exhaustive analysis of such potential transformation.

Like grid cells, border cellsalso have been reported from the MEC of bats during their crawling behavior (Yartsev et al., 2011). The question of the existence of these neurons in relation to complete 3D spatial navigation needs to be ascertained empirically. We predict the existence of neurons whose activity show bias towards any one or two walls of the box because the model neurons show this border like activity (Figures 6A-6D). Further analysis of border scores on orthogonal 2D planes proves that these neurons from the model are 3D extension of the previously reported 2D border cells in the hippocampal-parahippocampal regions (Bjerknes et al., 2014; Lever et al., 2009; Solstad et al., 2008). These cells can be termed as 3D border cells or wall cells. The other interesting aspect is that this neuronal cell type occupies a considerable amount of distribution, ~ 28%, in the LAHN (Figure 7C) which further reinforces the possibility of their existence.

LAHN neurons exhibit another interesting coding scheme by encoding a lower dimensional subspace i.e. a 2D plane while the animal performsa complete 3D navigation (Figures 6E-6G). We exclude border cells from this category even though border cells also code for a plane (walls of the box). This kind of lower dimensional coding has got important functional implications. For instance, as mentioned in the results section, a horizontal plane cell which has firing field arranged in a horizontal plane like fashion could potentially code for the altitude at which the animal is currently atand this information is critical for the efficacy of its navigation in higher dimensional space. This neuronal prediction could be empirically tested only when the animal engages inflight rather than crawl on the ground.

### Conclusion and future works

The proposed model explains the formation of 3D spatial representations using a hierarchical neural architecture comprising of both oscillatory and rate coded neurons. The model shows the essential learning rules that is required for the development of the 3D spatial maps using anti-hebbian neural network. The model captures the empirically reported 3D place cells and its isotropic nature (Ulanovsky, 2011; Yartsev and Ulanovsky, 2013; Yartsev et al., 2011). The model also captures the empirically reported anisotropy of grid cells during the animal’s navigation on a vertical plane (Hayman et al., 2011). It explains the dynamic nature of grid cell representations to account for a change in the trajectory statistics (variance of pitch distribution) of the animal. The model also sheds light on the possible planar symmetry exhibited by the grid cells in higher dimension under constrained pitch distribution. Apart from the aforementioned spatial cells, it predicts the possibility of two other types of spatial cells such as 3D border cells whose activity is biased to one or two walls of the environment and plane cells whose activity is restricted to a lower dimensional subspace i.e. a 2D plane in 3D space and such cells could possibly have implications in altitude coding.

Owing to the PCA like functionality of anti-hebbian network, the model is further analyzed and compared using PCA to get an intuitive understanding of the spatial cell formation in higher dimensions. Analysis using spatial cell descriptors such as spatial information index, grid score, border score and plane index shows an interesting formation of spatial bands on the eigen-spectrum. The place cell band occupies the high variance region, followed by the border cell band, and then comes the grid band and finally the plane cell band occupying the relatively low variance region when compared to the other aforementioned spatial cell types. Spatial cell distribution from the anti-hebbian network also follows this same order which points the possibility for the formation of spatial maps using PCA like unsupervised learning rule on the spatial information carrying signals like path integration signals.

To our knowledge, this is the first network level modeling effort to explain the formation of empirically reported 3D spatial cells along with the predictions on the other possible kinds of spatial cells. However, there are other possible add-ons to this network model. Firstly, the model is driven by the readily available direction information (both azimuth and pitch) which are assumed to come from the upstream areas like dorsal presubiclum (Finkelstein et al., 2015). This directional information could be extracted from the visual, proprioceptive and echolocation (in the case of bats) sensory information in a more biological way to produce a toroidal map of head direction using self-organization principles. This will further allow study of the influence of these sensory modalities on the 3D spatial cells. The next task is to analyze the representations of grid cells in higher dimensional space as the trajectory statistics change. What symmetry does it adopt when there isa change in the flight statistics of the animal? What are the possible dynamic configurations that it can assume to account for any change in the sensory stimulus? Thirdly, we would like to extend this model to study the goal directed navigation problem in 3D space where the animal searches for a target location and over the course of time it learns to find the target rapidly (this is a benchmark experimental paradigm used to study the 2D navigation problem under the name Morris water maze task (Vorhees and Williams, 2006)). This will shed light onto the possible way by which the 3D spatial cells aid the animal to find its goal location.

## MODEL EQUATIONS

### Azimuth and Pitch direction computation

Given change in the position across all the dimensions Δ_x_, Δ_y_, Δ_z_

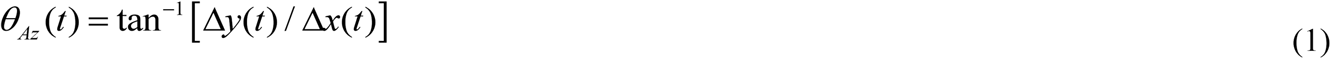

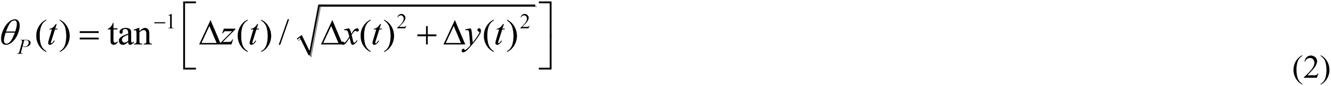

*θ_AZ_* and *θ_P_* are azimuth and pitch directions respectively.

### Head direction response

There are a total of *M* azimuth and pitch neurons divided in 7:3 ratio as per the empirical data(Finkelstein et al., 2015). Neurons in the azimuth and pitch are tuned to preferred directions that span 360°. Activity of neurons in each layer is computed as given below.

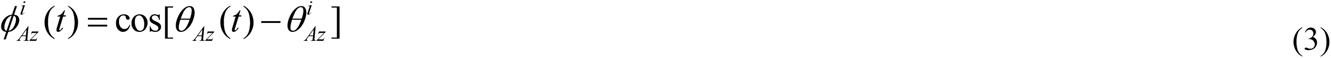

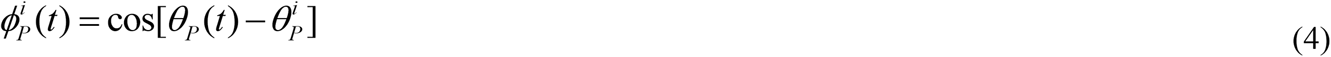

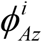 and 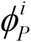 are the activities of the azimuth and pitch neurons respectively. 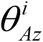 and 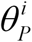 are the preferred directions of *i^th^* azimuth and pitch neurons respectively.

### Path integration layer

Each neuron in this layer is a phase oscillator whose dynamics is given in (5) and (6). This layer has one to-one connectivity with the preceding direction encoding layer. The direction and speed information is integrated into the phase of the oscillators.

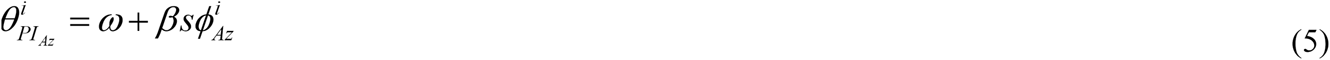

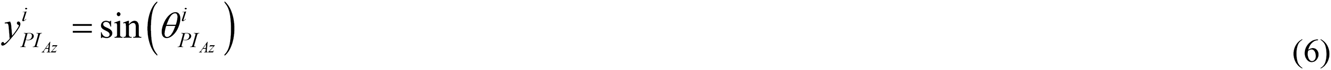

*ω* is the angular frequency of the oscillator such that *ω=2πfwhere f* is the base frequency of the oscillator. *β* is the modulation factor.

*s* is the speed of navigation given as

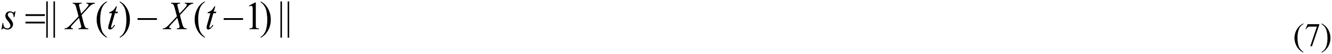

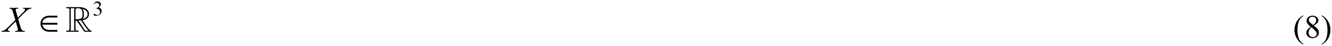

Empirically it is reported that the spectrum of LFP oscillations in the bat hippocampal area lack theta oscillations (Heys et al., 2013; Yartsev and Ulanovsky, 2013; Yartsev et al., 2011). Hence in the model also we used a very low base frequency *(f=* 0.5Hz) for the oscillators.

### Response equation of anti-hebbian network

The activity of a neuron in the Lateral Anti-Hebbian Network (LAHN) is given as (Földiak, 1990):

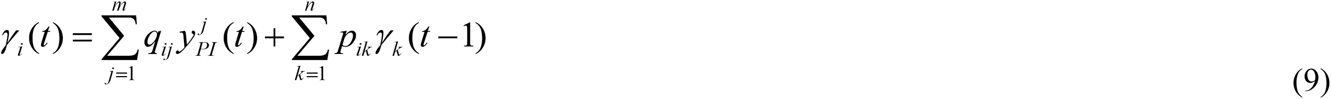

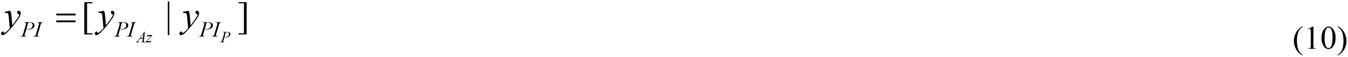

*γ_i_* is the activity of *i*^th^ LAHN neuron, *q_ij_* is the afferent weight connection from *j*^th^ component of input signal *y_PI_* to *i*^th^ neuron of LAHN, *p*_ik_ is the lateral weight connection from *k*^th^ to *i*^th^ LAHN neuron, *m* is the dimension of the input, and *n* is the total number of neurons in LAHN.

### Neural plasticity rule of anti-hebbian network

The plasticity rules for the synaptic weight connections of LAHN are as given below:

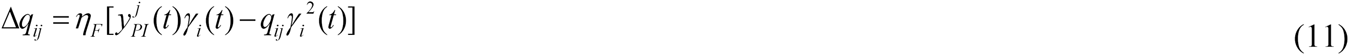

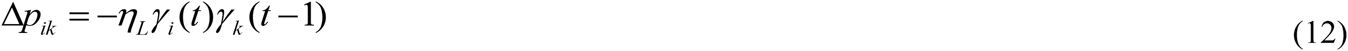

*Δq_ij_* is the change in the afferent weight connections (Hebbian rule)

*Δp_ij_* is the change in the lateral weight connections (Anti-hebbian rule)

*η*_F_and *η_L_* are the learning rates for the afferent and lateral weight connections respectively.

The information required for the unsupervised learning of LAHN neuron is available locally at its synaptic connections (Eqn (11) and (12)) and this makes the network biologically plausible (Földiak, 1990).

### 3D Firing rate map formation

3D volume of the physical space is binned into 41×41×41 voxels. Neuronal activity is assigned to the respective voxel depending on the firing field position. After this, the rate map is smoothed by a 3D Gaussian filter of *σ* = 3.

### 3D Autocorrelation map formation

Autocorrelation map is mainly computed to analyze the spatial periodicity of the neural activity by computing the respective gridness scores. Autocorrelation map, *r,* is computed as follows:

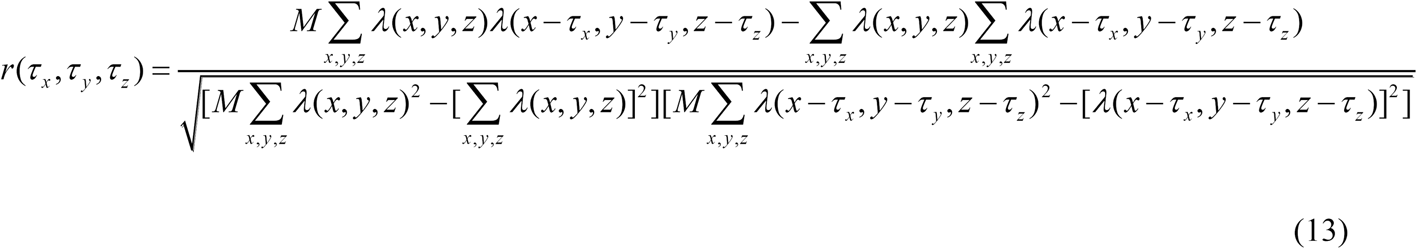

λ(x,y,z) is the firing rate at (x,y,z) location of the rate map, *M* is the total number of voxels in the rate map, *τ_x_, τ_y_* and *τ_z_* correspond to x, y and z coordinate spatial lags.

### Fitting a plane to the firing field

Since the firing field in this case has three dimensions, multiple regression is done to fit a plane to the firing field data. The model is as follows (Tabachnick and Fidell, 2007):

*ŷ= α_0_ + α_1_X_1_ + α_2_X_2_*

ŷ is the variable that has to be predicted.

*α* s are the regression coefficients that has to be determined

*x_1_* and *x_2_* are the regressors

Regressors are computed by minimizing the Sum of Squares of the Residuals (SSR) such as:

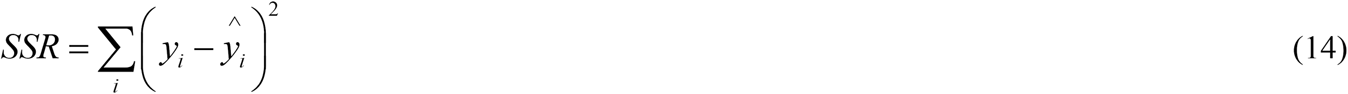

Y_i_ is the actual value and ŷ_I_ is the predicted value from the multiple regression model.

### Fitting an ellipsoid to the place cell firing field

Ellipsoid is fitted to the firing field data of a place cell by minimizing the following cost function.

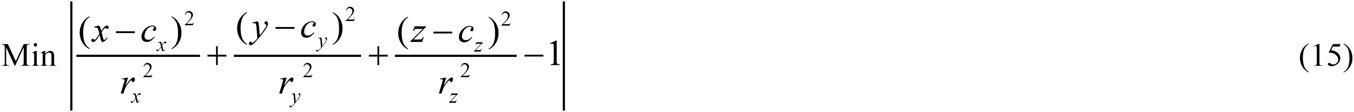

*C_x_*, *c_y_* and *c_z_* are the coordinates of the centre of the ellipsoid in the 3D Cartesian coordinate system. *r_x_, r_y_* and *r_z_* are the semi axis lengths of the ellipsoid.

### Computation of Spatial descriptors

1. **Spatial information index** This is used to classify if a neuron carries any spatially relevant information or not (Skaggs et al., 1993). It is computed from the neuron’s firing rate map as given below:

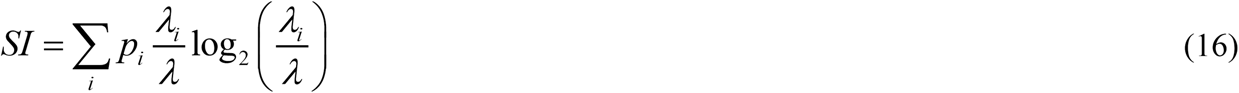

*p_i_* is the probability for the animal being in the z voxel. This can be computed as the ratio of the number of times the animal visited that voxel to the total time of its flight. *λ_ζ_* is the firing rate in the *i*^th^ voxel. *λ* is the mean firing rate across the rate map. SI is expressed as bits per spike.
2. **Grid score** Grid score is used to compute the rotational symmetry present in the autocorrelation map of the neuronal activity (Hafting et al., 2005). Given the autocorrelation map r, Hexagonal Gridness Score (HGS) is computed as follows:

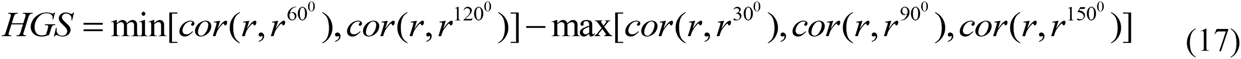

Square Gridness Score (SGS) is computed as follows (Soman et al., 2017):

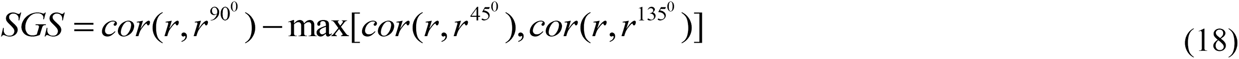

*r_θ_* is the autocorrelation map rotated by *θ*_°_
3. **Border score** Border score is used to analyze the border activity of a neuron. To compute border score, two quantities are assessed such as: the maximal extent of a single field on any wall *(CM)* and the mean firing distance *(d_M_). d_M_* is computed as the average distance of each bin in the firing rate map to the nearest wall, weighted by the firing rate activity in that bin. Border score (BS) is then computed as follows (Diehl et al., 2017; Solstad et al., 2008):

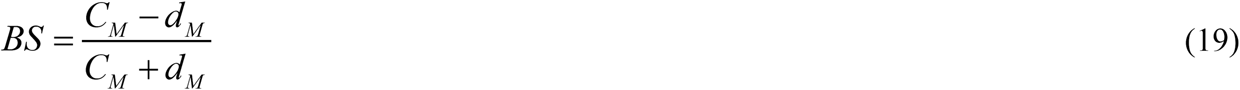

BS ranges between −1 and 1, where −1 represents a central firing and +1 represents a firing field that is perfectly aligned with a border.
4. **Plane index** Plane index is used to analyze the planeness of a neuron. This is obtained by fitting a plane to the 3D firing field, and the goodness of fit R^2^ value is considered as the plane index score. R^2^ value is the coefficient of determination whose value ranges between 0 and 1. It is defined as the proportion of the variance in the dependent variable that can be explained using the independent variable (Tabachnick and Fidell, 2007). R^2^ value can be estimated as follows:

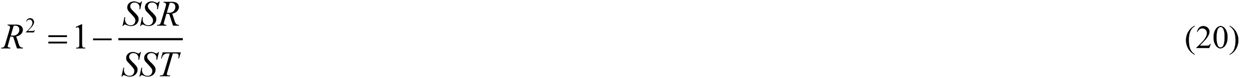

SSR is the sum of squared residuals as mentioned above. SST is the total sum of squares which gives the information on the total variance of the data around its mean. It is given as follows:

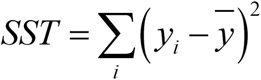 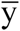 is the mean of the data to be predicted.

### Simulation parameters

Number of LAHN neurons = 50; *f* = 0.5 Hz; N_Az_ = 70; N_P_ = 30; *β* = 2; Spatial Information threshold = 1; Hexagonal Gridness Score threshold = 0; Border Score threshold = 0.5; Plain index threshold = 0.75; *η_F_* = 0.01; *η_L_* = 0.01;

Fitting of Gaussian distribution on the histogram of data is done using *histfit* function in MATLAB.

All simulations are done in MATLAB R2016a.

## References

Barlow, H., and Foldiak, P. (1989). The computing neuron. Adaptation and decorrelation in the cortex, 54–72.

Barry, C., Hayman, R., Burgess, N., and Jeffery, K.J. (2007). Experience-dependent rescaling of entorhinal grids. Nature neuroscience 10, 682–684.

Bassett, J.P., and Taube, J.S. (2001). Neural correlates for angular head velocity in the rat dorsal tegmental nucleus. Journal of Neuroscience 21, 5740–5751.

Bjerknes, T.L., Moser, E.I., and Moser, M.-B. (2014). Representation of geometric borders in the developing rat. Neuron 82, 71–78.

Bonnevie, T., Dunn, B., Fyhn, M., Hafting, T., Derdikman, D., Kubie, J.L., Roudi, Y., Moser, E.I., and Moser, M.-B. (2013). Grid cells require excitatory drive from the hippocampus. Nature neuroscience 16, 309–317.

Burak, Y., and Fiete, I.R. (2009). Accurate path integration in continuous attractor network models of grid cells. PLoS computational biology 5, e1000291.

Burgess, N., Barry, C., and O'keefe, J. (2007). An oscillatory interference model of grid cell firing. Hippocampus 17, 801–812.

Burgess, N., and O’Keefe, J. (2011). Models of place and grid cell firing and theta rhythmicity. Current opinion in neurobiology 21, 734–744.

Bush, D., and Burgess, N. (2014). A hybrid oscillatory interference/continuous attractor network model of grid cell firing. Journal of Neuroscience 34, 5065–5079.

Calton, J.L., and Taube, J.S. (2005). Degradation of head direction cell activity during inverted locomotion. Journal of Neuroscience 25, 2420–2428.

Conway, J.H., and Sloane, N.J.A. (2013). Sphere packings, lattices and groups, Vol 290 (Springer Science & Business Media).

Diehl, G.W., Hon, O.J., Leutgeb, S., and Leutgeb, J.K. (2017). Grid and nongrid cells in medial entorhinal cortex represent spatial location and environmental features with complementary coding schemes. Neuron 94, 83–92. e86.

Finkelstein, A., Derdikman, D., Rubin, A., Foerster, J.N., Las, L., and Ulanovsky, N. (2015). Threedimensional head-direction coding in the bat brain. Nature 517, 159–164.

Földiak, P. (1990). Forming sparse representations by local anti-Hebbian learning. Biological cybernetics 64, 165–170.

Fuhs, M.C., Redish, A.D., and Touretzky, D.S. (1998). A visually driven hippocampal place cell model. Computational neuroscience: Trends in research, 101–106.

Fuhs, M.C., and Touretzky, D.S. (2006). A spin glass model of path integration in rat medial entorhinal cortex. Journal of Neuroscience 26, 4266–4276.

Geva-Sagiv, M., Romani, S., Las, L., and Ulanovsky, N. (2016). Hippocampal global remapping for different sensory modalities in flying bats. Nature neuroscience 19, 952–958.

Hafting, T., Fyhn, M., Molden, S., Moser, M.-B., and Moser, E.I. (2005). Microstructure of a spatial map in the entorhinal cortex. Nature 436, 801–806.

Hayman, R., Verriotis, M.A., Jovalekic, A., Fenton, A.A., and Jeffery, K.J. (2011). Anisotropic encoding of three-dimensional space by place cells and grid cells. Nature neuroscience 14, 1182–1188.

Hayman, R.M., Casali, G., Wilson, J.J., and Jeffery, K.J. (2015). Grid cells on steeply sloping terrain: evidence for planar rather than volumetric encoding. Frontiers in psychology 6.

Heys, J.G., MacLeod, K.M., Moss, C.F., and Hasselmo, M.E. (2013). Bat and rat neurons differ in theta frequency resonance despite similar coding of space. Science 340, 363–367.

Knierim, J.J., and McNaughton, B.L. (2001). Hippocampal place-cell firing during movement in threedimensional space. Journal of Neurophysiology 85, 105–116.

Kropff, E., and Treves, A. (2008). The emergence of grid cells: Intelligent design or just adaptation? Hippocampus 18, 1256–1269.

Lever, C., Burton, S., Jeewajee, A., O'Keefe, J., and Burgess, N. (2009). Boundary vector cells in the subiculum of the hippocampal formation. Journal of Neuroscience 29, 9771–9777.

Mathis, A., Stemmler, M.B., and Herz, A.V. (2015). Probable nature of higher-dimensional symmetries underlying mammalian grid-cell activity patterns. Elife 4, e05979.

Moser, E.I., Roudi, Y., Witter, M.P., Kentros, C., Bonhoeffer, T., and Moser, M.-B. (2014). Grid cells and cortical representation. Nature Reviews Neuroscience 15, 466–481.

O'Keefe, J., and Dostrovsky, J. (1971). The hippocampus as a spatial map. Preliminary evidence from unit activity in the freely-moving rat. Brain research 34, 171–175.

Oja, E. (1982). Simplified neuron model as a principal component analyzer. Journal of mathematical biology 15, 267–273.

Omer, D.B., Maimon, S.R., Las, L., and Ulanovsky, N. (2018). Social place-cells in the bat hippocampus. Science 359, 218–224.

Sanger, T.D. (1989). Optimal unsupervised learning in a single-layer linear feedforward neural network. Neural Networks 2, 459–473.

Skaggs, W.E., McNaughton, B.L., and Gothard, K.M. (1993). An information-theoretic approach to deciphering the hippocampal code. In Advances in neural information processing systems, pp. 1030–1037.

Solstad, T., Boccara, C.N., Kropff, E., Moser, M.-B., and Moser, E.I. (2008). Representation of geometric borders in the entorhinal cortex. Science 322, 1865–1868.

Soman, K., Muralidharan, V., and Chakravarthy, V.S. (2017). A Model of Multisensory Integration and its Influence on Hippocampal Spatial Cell Responses. IEEE Transactions on Cognitive and Developmental Systems, In Press.

Stackman, R.W., and Taube, J.S. (1998). Firing properties of rat lateral mammillary single units: head direction, head pitch, and angular head velocity. Journal of Neuroscience 18, 9020–9037.

Stackman, R.W., Tullman, M.L., and Taube, J.S. (2000). Maintenance of rat head direction cell firing during locomotion in the vertical plane. Journal of Neurophysiology 83, 393–405.

Stella, F., and Treves, A. (2015). The self-organization of grid cells in 3D. Elife 4, e05913.

Tabachnick, B.G., and Fidell, L.S. (2007). Using multivariate statistics (Allyn & Bacon/Pearson Education).

Taube, J.S., and Bassett, J.P. (2003). Persistent neural activity in head direction cells. Cerebral Cortex 13, 1162–1172.

Taube, J.S., Muller, R.U., and Ranck, J.B. (1990a). Head-direction cells recorded from the postsubiculum in freely moving rats. I. Description and quantitative analysis. Journal of Neuroscience 10, 420–435.

Taube, J.S., Muller, R.U., and Ranck, J.B. (1990b). Head-direction cells recorded from the postsubiculum in freely moving rats. II. Effects of environmental manipulations. Journal of Neuroscience 10, 436–447.

Ulanovsky, N. (2011). Neuroscience: how is three-dimensional space encoded in the brain? Current Biology 21, R886–R888.

Ulanovsky, N., and Moss, C.F. (2007). Hippocampal cellular and network activity in freely moving echolocating bats. Nature neuroscience 10, 224.

Vorhees, C.V., and Williams, M.T. (2006). Morris water maze: procedures for assessing spatial and related forms of learning and memory. Nature protocols 1, 848.

Widloski, J., and Fiete, I.R. (2014). A model of grid cell development through spatial exploration and spike time-dependent plasticity. Neuron 83, 481–495.

Wold, S., Esbensen, K., and Geladi, P. (1987). Principal component analysis. Chemometrics and intelligent laboratory systems 2, 37–52.

Yartsev, M.M., and Ulanovsky, N. (2013). Representation of three-dimensional space in the hippocampus of flying bats. Science 340, 367–372.

Yartsev, M.M., Witter, M.P., and Ulanovsky, N. (2011). Grid cells without theta oscillations in the entorhinal cortex of bats. Nature 479, 103–107.

Zilli, E.A., and Hasselmo, M.E. (2010). Coupled noisy spiking neurons as velocity-controlled oscillators in a model of grid cell spatial firing. Journal of Neuroscience 30, 13850–13860.

